# *De novo* mutations in the GTP/GDP-binding region of RALA, a RAS-like small GTPase, cause intellectual disability and developmental delay

**DOI:** 10.1101/378349

**Authors:** Susan M. Hiatt, Matthew B. Neu, Ryne C. Ramaker, Andrew A. Hardigan, Jeremy W. Prokop, Miroslava Hancarova, Darina Prchalova, Marketa Havlovicova, Jan Prchal, Viktor Stranecky, Dwight K.C. Yim, Zöe Powis, Boris Keren, Caroline Nava, Cyril Mignot, Marlene Rio, Anya Revah-Politi, Parisa Hemati, Nicholas Stong, Alejandro D. Iglesias, Sharon F. Suchy, Rebecca Willaert, Ingrid M. Wentzensen, Patricia G. Wheeler, Lauren Brick, Mariya Kozenko, Anna C.E. Hurst, James W. Wheless, Yves Lacassie, Richard M. Myers, Gregory S. Barsh, Zdenek Sedlacek, Gregory M. Cooper

## Abstract

Mutations that alter signaling of RAS/MAPK-family proteins give rise to a group of Mendelian diseases known as RASopathies, but the matrix of genotype-phenotype relationships is still incomplete, in part because there are many RAS-related proteins, and in part because the phenotypic consequences may be variable and/or pleiotropic. Here, we describe a cohort of ten cases, drawn from six clinical sites and over 16,000 sequenced probands, with *de novo* protein-altering variation in *RALA*, a RAS-like small GTPase. All probands present with speech and motor delays, and most have intellectual disability, low weight, short stature, and facial dysmorphism. The observed rate of *de novo RALA* variants in affected probands is significantly higher (p=4.93 × 10^−11^) than expected from the estimated mutation rate. Further, all *de novo* variants described here affect conserved residues within the GTP/GDP-binding region of *RALA*; in fact, six alleles arose at only two codons, Val25 and Lys128. We directly assayed GTP hydrolysis and RALA effector-protein binding, and all but one tested variant significantly reduced both activities. The one exception, S157A, reduced GTP hydrolysis but significantly increased RALA-effector binding, an observation similar to that seen for oncogenic RAS variants. These results show the power of data sharing for the interpretation and analysis of rare variation, expand the spectrum of molecular causes of developmental disability to include *RALA*, and provide additional insight into the pathogenesis of human disease caused by mutations in small GTPases.

**Author Summary:** While many causes of developmental disabilities have been identified, a large number of affected children cannot be diagnosed despite extensive medical testing. Previously unknown genetic factors are likely to be the culprits in many of these cases. Using DNA sequencing, and by sharing information among many doctors and researchers, we have identified a set of individuals with developmental problems who all have changes to the same gene, *RALA.* The affected individuals all have similar symptoms, including intellectual disability, speech delay (or no speech), and problems with motor skills like walking. In nearly all of these cases (10 of 11), the genetic change found in the child was not inherited from either parent. The locations and biological properties of these changes suggest that they are likely to disrupt the normal functions of RALA and cause significant health problems. We also performed experiments to show that the genetic changes found in these individuals alter two key functions of RALA. Together, we have provided evidence that genetic changes in *RALA* can cause DD/ID. These results will allow doctors and researchers to identify additional children with the same condition, providing a clinical diagnosis to these families and leading to new research opportunities.

## Introduction

Developmental delay and intellectual disability (DD/ID) affect about 1-2% of individuals worldwide [1]. Many highly penetrant genetic variants underlying DD/ID have been identified, but a large fraction of disease risk remains unexplained [2, 3]. While some DD/ID-cases may result from environmental factors and small-effect common variants [4] it is likely that many probands harbor pathogenic, highly penetrant variation in as-yet-unknown disease-associated genes.

The RASopathies are a group of genetic conditions often associated with developmental disorders [5], having in common mutational disruption of genes in the RAS/MAPK pathway that alter patterns of signal transduction. RASopathies are individually rare and pleiotropic but are collectively one of the most common causes of developmental disorders. Associated features include neurocognitive impairment, craniofacial dysmorphology, anomalies of the cardiovascular and musculoskeletal systems, cutaneous lesions, and increased risk of tumor formation [6]. For example, variation in *HRAS* is associated with Costello Syndrome (MIM:218040) and Noonan Syndrome (MIM:609942), variation in *KRAS* is associated with Cardiofaciocutaneous syndromes (MIM:615278), and variation in *NRAS* has been observed in probands with RASopathy-associated phenotypes [7].

Given the genetic and phenotypic heterogeneity among DD/ID in general and RASopathies in particular, collaboration and data sharing among clinicians, researchers, and sequencing centers is necessary to enable, or accelerate, discoveries of new forms of disease. One tool to facilitate such collaborations is GeneMatcher, launched in 2013 as a way to connect researchers and clinicians with interests in specific genes [8].

Here, we present details of a cohort, assembled via GeneMatcher, of eleven total probands (including one set of monozygotic twins) with protein-altering variation in *RALA*, which encodes a RAS-like small GTPase; the variants arose *de novo* in ten of these probands. All probands present with developmental delay. Detailed phenotyping, computational analyses of observed variation, and functional studies lead to the conclusion that variation affecting the GTPase activity and downstream signaling of RALA underlies a new neurodevelopmental RASopathy-like disorder.

## Results

This study originated as a collaboration facilitated by GeneMatcher through shared interests in *RALA* as a result of observations from exome sequencing (ES) or genome sequencing (GS) as part of DD/ID-related clinical or research testing. In the Methods and Appendix S1, we describe the research sites that identified one or more affected probands reported in this study, the methods used for sequencing and analysis, and related details. In total, we identified *RALA* mutations in eleven affected probands from ten unrelated families. These variants were identified from a combined cohort of over 16,000 probands sequenced by six groups who independently submitted *RALA* to GeneMatcher (Table 1, Appendix S1).

**Table 1.**
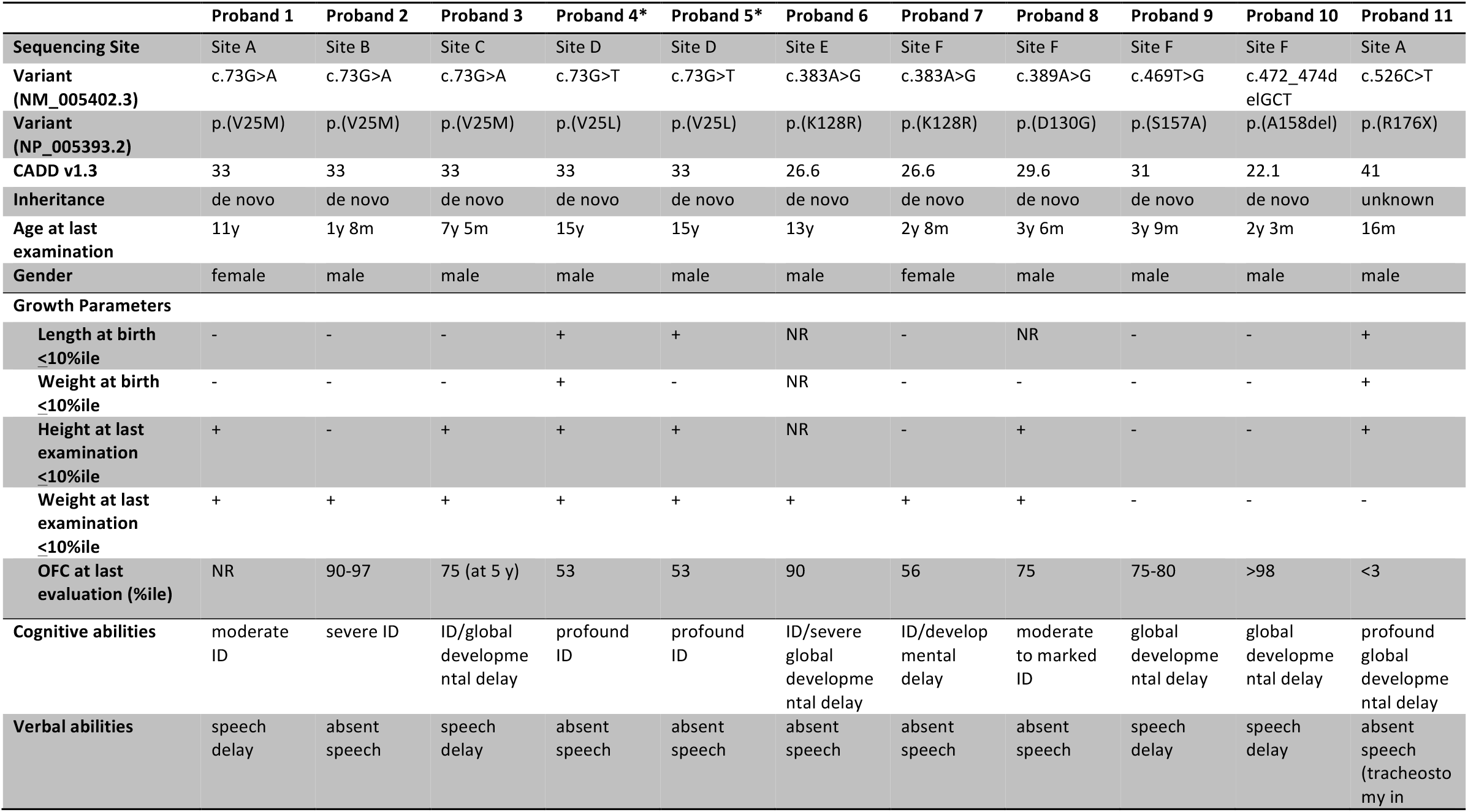
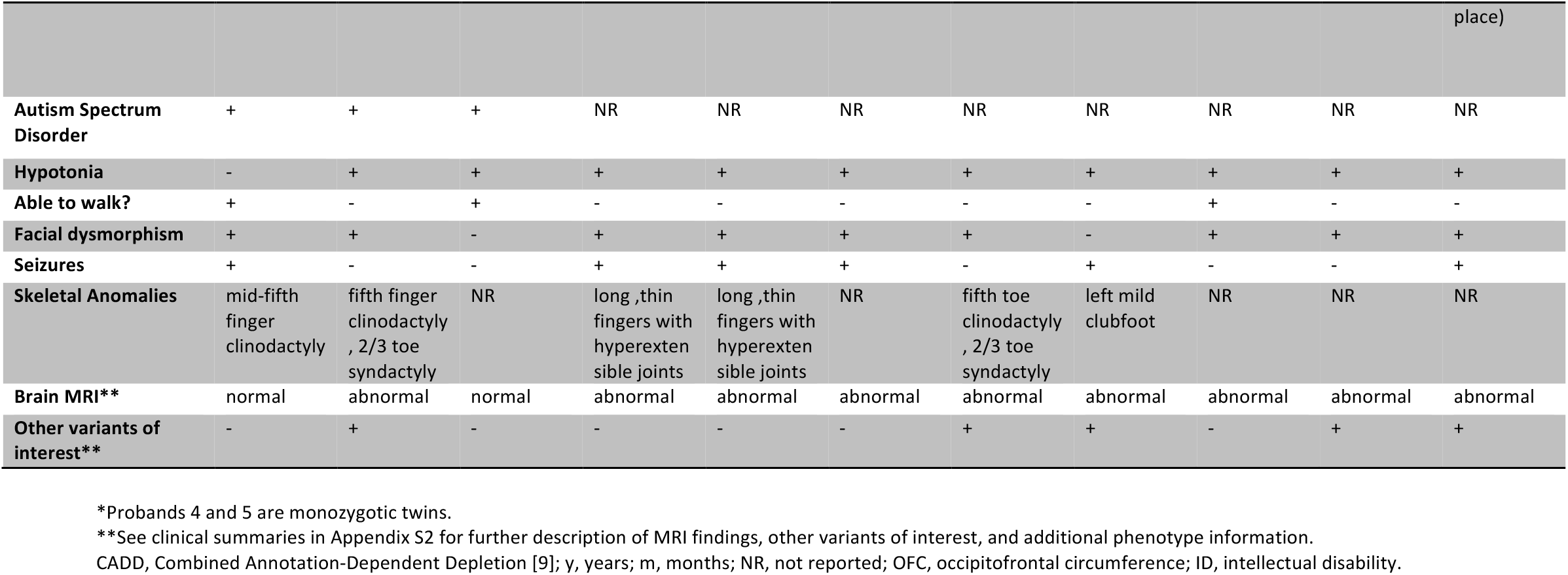
Genotypes and phenotypes of individuals with variation in *RALA.*

### Phenotypic details

All eleven probands presented with speech problems, including absent speech in seven and speech delay in the remaining four. Ten of the eleven probands are reported to have hypotonia, with eight unable to walk. Intellectual disability was specifically noted for 8 of 11, (but not ruled out for the remaining three, see Table 1). Birth measurements were available for nine probands and three (33%) reported either length or weight (or both) at less than the tenth percentile. Height and weight measurements at last examination were available for all probands (except for height in one). Six of ten probands (60%) were reported to have heights less than the 10th percentile at last examination, while eight of eleven (73%) were reported to have weights less than the 10^th^ percentile. Three probands had head circumference measurements greater than or equal to the 90^th^ percentile at last evaluation. Nine of eleven probands were reported to have dysmorphic facial features. Several consistent features were observed, including a broad, prominent forehead, horizontal eyebrows, epicanthus, mild ptosis, slightly anteverted nares, wide nasal bridge, short philtrum, thin upper lip vermillion with an exaggerated Cupid’s bow, pointed chin, and low-set ears with increased posterior angulation (Figure 1).

Additional common but variable features were observed: seizures were present in most probands (6/11), as were structural brain abnormalities detected by MRI (9/11). Six of eleven probands were reported to have skeletal anomalies such as clinodactyly (3 of 6) and/or 2/3 toe syndactyly (2 of 6). None of the probands are reported to have had cancer. Clinical summaries with additional details are available in Appendix S2.

### Molecular characterization of variation

Genetic variation within this cohort includes eight *de novo* heterozygous missense variants (in nine probands, including the monozygotic twin pair), one *de novo* heterozygous in-frame deletion of one amino acid, and one heterozygous premature stop of unknown inheritance (Table 1, Figure 2A). Except for R176X (see below), all observed variants are absent from gnomAD [10] and TopMed genomes (“Bravo”) [11]. These variants have CADD scores ranging from 22.1 to 41 suggesting they are highly deleterious, similar to the majority of mutations previously reported to cause Mendelian diseases [9].

**Figure 2.**
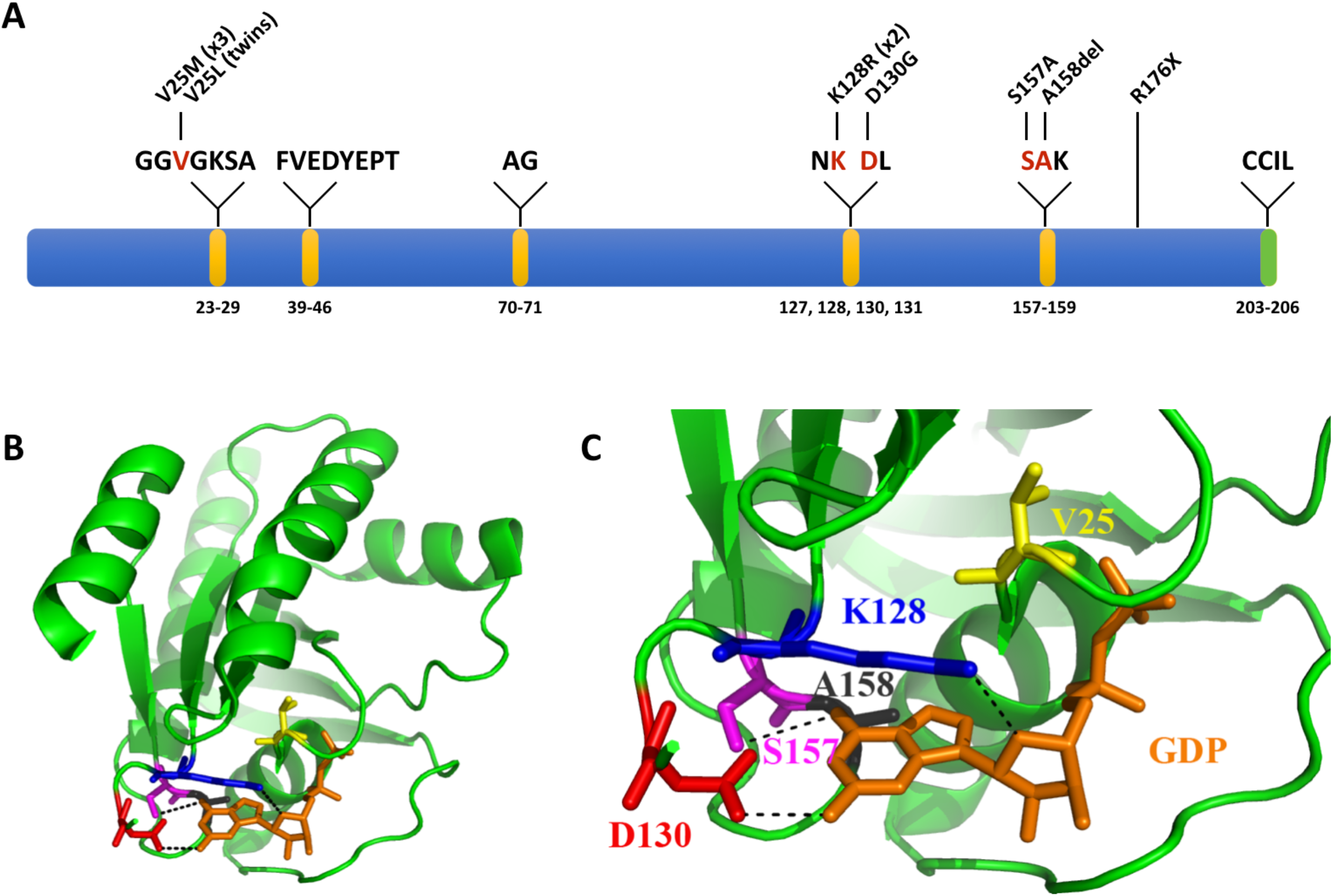
Variation observed in RALA clusters in GTP/GDP-binding regions. A. Linear model of RALA, including GTP/GDP-binding regions (depicted in yellow, as defined by molecular modeling data) and the CAAX motif (CCIL in the case of RALA; depicted in green). Positions of amino acid residues that form the GTP/GDP-binding region are listed below the model, and residues within those regions are listed above the model. Residues affected by variation observed here are shown in red. The predicted protein changes for described variation are shown above the affected amino acid residues. B. Positions of RALA amino acid residues affected by variation relative to the GDP molecule. C. A zoomed in view of the variation observed within the GTP/GDP-binding region. GDP is shown in a licorice representation in orange. The RALA protein is shown in a cartoon representation in green, with the mutated residues in licorice representation. V25 is in yellow, K128 in blue, D130 in red, S157 in magenta, and A158 in black. Hydrogen bonds between the side chains of these amino acids and GDP are shown as black dashed lines. See Supplemental Figures S1-S5, S8 for consequences of individual variants on the protein structure.

Seven probands (1-7), including the monozygotic twin pair, harbor recurrent *de novo* variants affecting one of only two codons, those encoding residues Val25 and Lys128, while the remaining three *de novo* variants affect Asp130, Ser157, and Ala158. All of these residues are computationally annotated as one of 24 residues, within a total protein length of 206 amino acids, that form the GTP/GDP-binding region of the RALA protein (Figure 2, Methods). While Val25 does not directly interact with GTP/GDP, variation observed at this position (Val25Met and Val25Leu) would likely result in distortion of the structure of the GTP/GDP-binding pocket (Figure 2B, 2C, Supplemental Figure S1). Lys128, Asp130 and Ser157 all form hydrogen bonds with GTP/GDP in the wild type protein (Figure 2B, 2C, Supplemental Figures S2-S4). Although Lys128Arg would retain the positive charge of the side chain, steric hindrance resulting from the larger size of the Arg side chain would likely result in disruption of this binding pocket (Supplemental Figure S2). Both Asp130Gly and Ser157Ala are predicted to result in loss of hydrogen bond formation (Figure 2B, 2C, Supplemental Figures S3, S4). The remaining *de novo* variant, an in-frame deletion of Ala158, results in a shift of Lys159 into the GTP/GDP binding region of RALA, which likely hinders GTP/GDP binding (Supplemental Figure S5). Variation at all five of these residues is thus predicted to alter GTP/GDP binding. This conclusion is consistent with the high degree of conservation at these residues throughout evolution in RALA (Supplemental Figure S6) as well as in other related genes including HRAS, KRAS, and NRAS (Supplemental Figure S7) and RAP1A/B and RHOA[12].

The predicted nonsense variant Arg176X in proband 11 lies within the last exon of *RALA*, and thus may not result in nonsense-mediated decay (NMD) of the transcript. This would yield a protein that lacks the 29 C-terminal residues (Supplemental Figure S8), which are known to contain at least two critical regulatory regions. Phosphorylation of Ser194 by Aurora kinase A (AURKA) activates RALA, affects its localization, and results in activation of downstream effectors like RALBP1 [13, 14]. Additionally, the C-terminal CAAX motif (CCIL in the case of RALA) is essential for proper localization and activation of RALA via prenylation of Cys203 [15, 16].

### Enrichment and clustering of missense variation

We next assessed whether the *de novo* variants in our cohort were enriched compared to that which would be expected in the absence of a disease association. Eight unrelated individuals were drawn from cohorts of at least 400 proband-parent trios, collectively spanning 16,086 probands (Appendix S1). When comparing the frequency of observed *de novo* variation to the expected background frequency of *de novo* missense or loss-of-function variation in *RALA* (6.16 × 10^−6^ per chromosome) [17], we find a highly significant enrichment for *de novo* variants in affected probands (8 observed *de novo* variants in 32172 screened alleles vs. 0.198 expected, Exact Binomial test p=4.93 × 10^−11^). We note that this p-value is likely conservative, as it results from comparison of the observed rate to the expected frequency of *de novo* variation over the entire gene. However, six of the nine *de novo* alleles affect only two codons, and all observed *de novo* variants are within the GTP-interacting space of 24 residues (11.7% of the 206-aa protein, Figure 2A). This clustering likely reflects a mechanism of disease that depends specifically on alterations to GTP/GDP binding and, subsequently, RALA signaling.

Population genetic data also support pathogenicity of these variants. *RALA* has a pLI score of 0.95 in ExAC [10], suggesting that it is intolerant to loss-of-function variation. While *RALA* has an RVIS score rank [18] of 50.45%, it also has an observed/expected ratio percentile of 0.92%, a score that has been suggested to be more accurate for small proteins wherein observed and expected allele counts are relatively small [19]. Furthermore, population genetic data also support the likely special relevance of mutations in the GTP/GDP-binding pocket. No high-quality (“PASS” only) missense variants are observed at any frequency at any of the 24 GTP/GDP-coordinating residues in either gnomAD [10] or BRAVO[11]; in contrast, there are missense variants observed at 34 of the 182 RALA residues outside the GTP/GDP-interaction region (Supplemental Table S1). This distribution across *RALA* is likely non-random (Fisher’s exact test p=0.017) and suggestive of especially high variation intolerance in this region of RALA.

### Comparison to disease associated with RAS-family GTPases

RALA and other RAS-family GTPases have a high degree of similarity, and germline variation in other RAS-family GTPases is known to be associated with developmental disorders [5]. Comparisons of phenotypes observed here to those reported in these RASopathies suggest considerable overlap, including DD/ID, growth retardation, macrocephaly, high broad forehead and mildly dysplastic dorsally rotated ears. Further, we compared the specific variants observed here to variants in HRAS, KRAS, or NRAS, reported as pathogenic for RASopathies (Supplemental Table S2, Supplemental Figure S7). *De novo* heterozygous missense variation at Val14 of KRAS, the homologous equivalent of Val25 in RALA, was previously reported in four unrelated individuals with Noonan syndrome [20, 21]. Functional studies showed that this variant may alter intrinsic and stimulated GTPase activity and may increase the rate of GDP release [20, 21]. A *de novo* variant in HRAS at Lys117, the homologous equivalent of Lys128 in RALA, was found in two unrelated probands with Costello Syndrome [22]. Lastly, a *de novo* HRAS variant at Ala146, the homologous equivalent of Ala158 in RALA, was reported in at least three patients with Costello Syndrome [23]. Variation at this residue has also been reported as a recurrent somatic variant in colorectal cancers [24].

### Functional analysis

We investigated the functional consequences of the variants described above by expressing and purifying recombinant RALA proteins, and then measuring their abilities to hydrolyze GTP and to interact with an immobilized RALA effector protein (see Methods). While wild-type RALA showed robust GTPase activity under these experimental conditions, all mutants tested here exhibited a dramatic reduction in GTPase activity, including a mutant RALA that was not observed in probands but which carries a missense substitution, G23D, homologous to the G12D KRAS or HRAS variant commonly observed in tumor tissue (Figure 3A).

**Figure 3.**
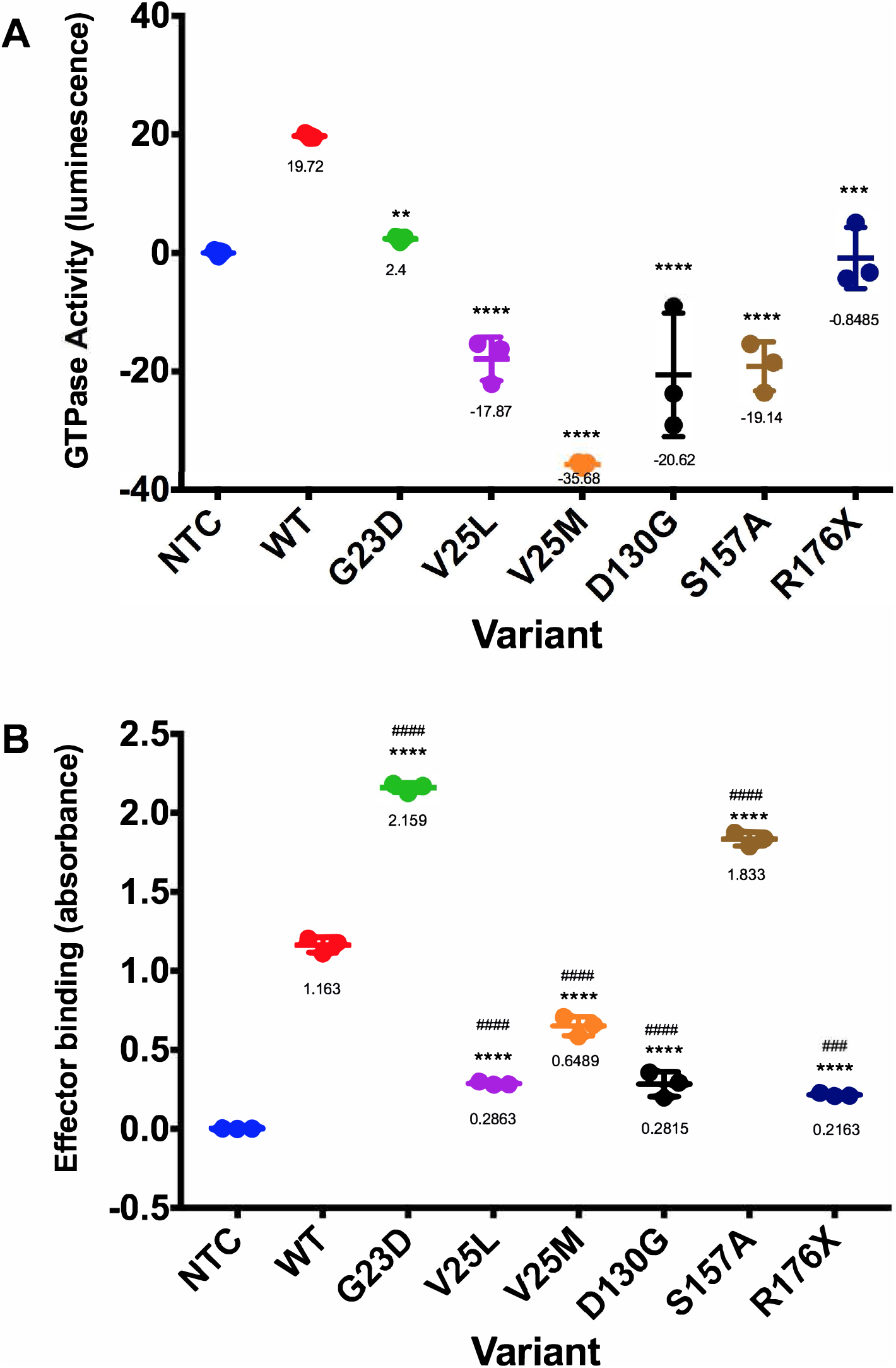
Missense variation in RALA affects GTPase activity and RALA effector binding. A. GTPase activity of purified recombinant RALA proteins was assessed using a luminescence assay. Raw luminescence values (measuring remaining free GTP) were subtracted from 100 to calculate activity, and were then normalized to a no template control (NTC). WT, wild-type RALA. G23D, predicted constitutively active mutant (not from a proband). ** indicates p-value = 0.0015 compared to WT, *** indicates p-value = 0.0003, and **** indicates p-value < 0.0001 compared to WT. Mean values of one experiment performed in triplicate are shown. B. Binding of purified recombinant RALA proteins to an effector molecule was assessed using an ELISA-based assay. Absorbances were normalized to a no template control (NTC). Mean values of one experiment performed in triplicate are shown. WT, wild-type RALA. **** indicates p-value < 0. 0001 compared to WT. #### indicates p-value < 0.0001 compared to NTC. ### indicates p-value = 0.0001 compared to NTC.

As GTPase activity of mutant RAS family proteins alone is not always a clear indication of downstream effects [20, 21], we also assessed binding of these mutants to a RALA effector protein using an ELISA-based method (see Methods). In this assay, recombinant G23D RALA protein exhibited approximately two-fold increased binding (p < 0.0001, Figure 3B), as anticipated for a constitutively active gain-of-function alteration [20, 25]. V25L, V25M, D130G and R176X each showed a roughly 2-5-fold reduction in effector binding compared to wild-type (each p < 0.0001, Figure 3B). In contrast, the S157A mutant exhibited increased binding compared to wild-type, suggesting that it may act in a constitutively-active manner similar to G23D (p < 0.0001, Figure 3B). We note that while there is some variation among mutants in the efficiency of protein production and purification (Methods, Supplemental Figure S9), whether or not one normalizes to relative band intensity from Western blots of purified protein does not qualitatively affect these conclusions (Supplemental Figure S10).

### Other candidate variants in probands with RALA variants

In this and other cases of rare disease sequencing, it is important to consider other variation in any given patient that may be pathogenic. In six of the eleven cases presented here, the *RALA* variant was found to be the only plausible candidate. In five cases, other variants were discovered that were also initially considered as potential disease-causing mutations. (Table 1, Appendix S2). Proband 2 has a hemizygous variant in *FLNA* (p.V606L), inherited from his unaffected heterozygous mother. Phenotype comparison, consultation with a filaminopathy disease expert, and application of the ACMG variant interpretation guidelines [26] resulted in the scoring of this variant as a VUS. Interestingly, FLNA is an interaction partner of RALA [27], but the relevance of this variant is unclear. Proband 7 has a *de novo* variant in *SHANK2* (p.A1101T); however, this allele is present in gnomAD three times and thus is not likely to be a highly penetrant allele resulting in DD/ID. Proband 8 has a variant in *SCN1A* p.R187Q; however, this variant was inherited from an unaffected father, is present in gnomAD in one heterozygote, and, according to the referring clinician, the phenotype observed in the proband is not consistent with Dravet syndrome. Finally, proband 10 carries a paternally-inherited 1.349 Mb duplication of 1q21.1-q21.2. This duplication has been reported to be associated with mild to moderate DD/ID, autism spectrum disorders, ADHD and behavioral problems, and other variable features [28]. While the patient may have some phenotypic features of this duplication, the patient’s MRI findings and severity of delays are not likely explained by this inherited duplication.

Proband 11 carries a nonsense variant, R176X, which is unusual given the apparent specificity for the GTP/GDP-binding region of RALA observed in the other cases in our cohort. Clinically, we consider the R176X to be a variant of uncertain significance for several reasons. The R176X allele has been observed twice in the Bravo genome database, and parental DNA for this proband was not available, so we do not know whether the variant is *de novo* or inherited. In addition, the proband has microcephaly and more profound delays than others in the cohort, and also has large regions of homozygosity consistent with parental consanguinity. These regions of homozygosity suggests an additional and/or more complex molecular pathogenesis.

## Discussion

The rapidly expanding application of genome sequencing to clinical settings is rapidly expanding our knowledge of mutations that cause rare disease, and has engendered new strategies for analysis, new rubrics for molecular pathology, and new platforms for collaboration. Here we apply these advances to show that mutations in the GTP/GDP-binding region of *RALA* cause developmental and speech delay, together with minor dysmorphic features. Mutations in RAS family members and RAS signaling pathways are well-recognized causes of several dysmorphic syndromes and cancer, but germline mutations in *RALA* have not been previously associated with disease. Our results add to basic knowledge about the biology and function of RAS family members, raise new questions about the molecular pathogenesis of mutations that affect small GTPases, and have important implications for clinical genomics.

Among the small GTPases, RALA and RALB are the most closely related to the RAS subfamily (~50% amino acid similarity), and function as a third arm of the RAS effector pathway in addition to RAF and PI3K activation [5]. RALA and RALB have different expression patterns— RALA is broadly expressed whereas expression of RALB is enriched in endocrine tissues [29] — but also exhibit some degree of genetic redundancy: in gene-targeted mice, loss of function for RALA causes a severe neural tube defect that is exacerbated by simultaneous loss of RALB [30]. In neuronal culture systems, RALA has been implicated in the development, plasticity, polarization, migration, branching, and spine growth of neurons [31–35], as well as the renewal of synaptic vesicles and trafficking of NMDA, AMPA, and dopamine receptors to the postsynaptic membrane [27, 34, 36]. These studies evaluated the effects of RALA in multiple ways, including through loss of function studies (e.g., using RNA interference), and designed mutational alterations to GTP/GDP hydrolysis, suggesting that multiple types of RALA perturbation have molecular and cellular consequences. Several aspects of our results suggest that developmental delay in humans is not caused by a simple loss-of-function for RALA. Our patients are all heterozygous, whereas in mice, heterozygosity for loss of function does not obviously affect development or viability [30]. In our functional assay, all of the proband alleles exhibited reduced GTPase activity, similar to most oncogenic RAS alleles. However, they exhibited variability in their ability to bind RALA effector protein, with one showing increased effector binding and the others all reducing effector binding. Simillar variability of *in vitro* functional effects were reported for *KRAS* GTP/GDP-binding domain mutations observed in patients with developmental disorders [21]. Importantly, all of the proband alleles assessed here to be pathogenic are *de novo* missense variants in the GDP/GTP-binding domain, including six recurring at only two codons. This observation is in contrast to the variation in large human population datasets which is only observed outside of this domain. Taken together, our data suggest that the molecular pathogenesis of developmental delay in the patients described here is brought about by a genetic mechanism that specifically depends on perturbations to the normal GTP-GDP cycling of RALA.

In summary, we show that *de novo* variation affecting *RALA* in individuals with DD/ID is highly enriched compared to background mutational models, exhibits clear spatial clustering in/near to the GTP/GDP-binding region, tends to affect positions whose homologous equivalents in other small GTPases are reported to harbor disease-associated variation, and significantly alters GTPase activity and RALA effector binding in *in vitro* functional assays. These observations add to the diverse and pleiotriopic group of Mendelian disorders caused by variation in RAS-family GTPases and related RAS pathways.

## Materials and Methods

### Informed consent

Informed consent to publish de-identified data was obtained from all participating families, and informed consent to publish clinical photographs was also obtained when applicable. Collection and analysis of sequencing data from all participants was conducted with the approval of appropriate human subjects research governing bodies.

### Exome/Genome sequencing

Exome sequencing (ES) or genome sequencing (GS) was performed at each of the following sites in either a research or clinical setting. Additional details, including cohort sizes used in p-value calculations, are provided in Supplemental Materials and Methods, Appendix S1.

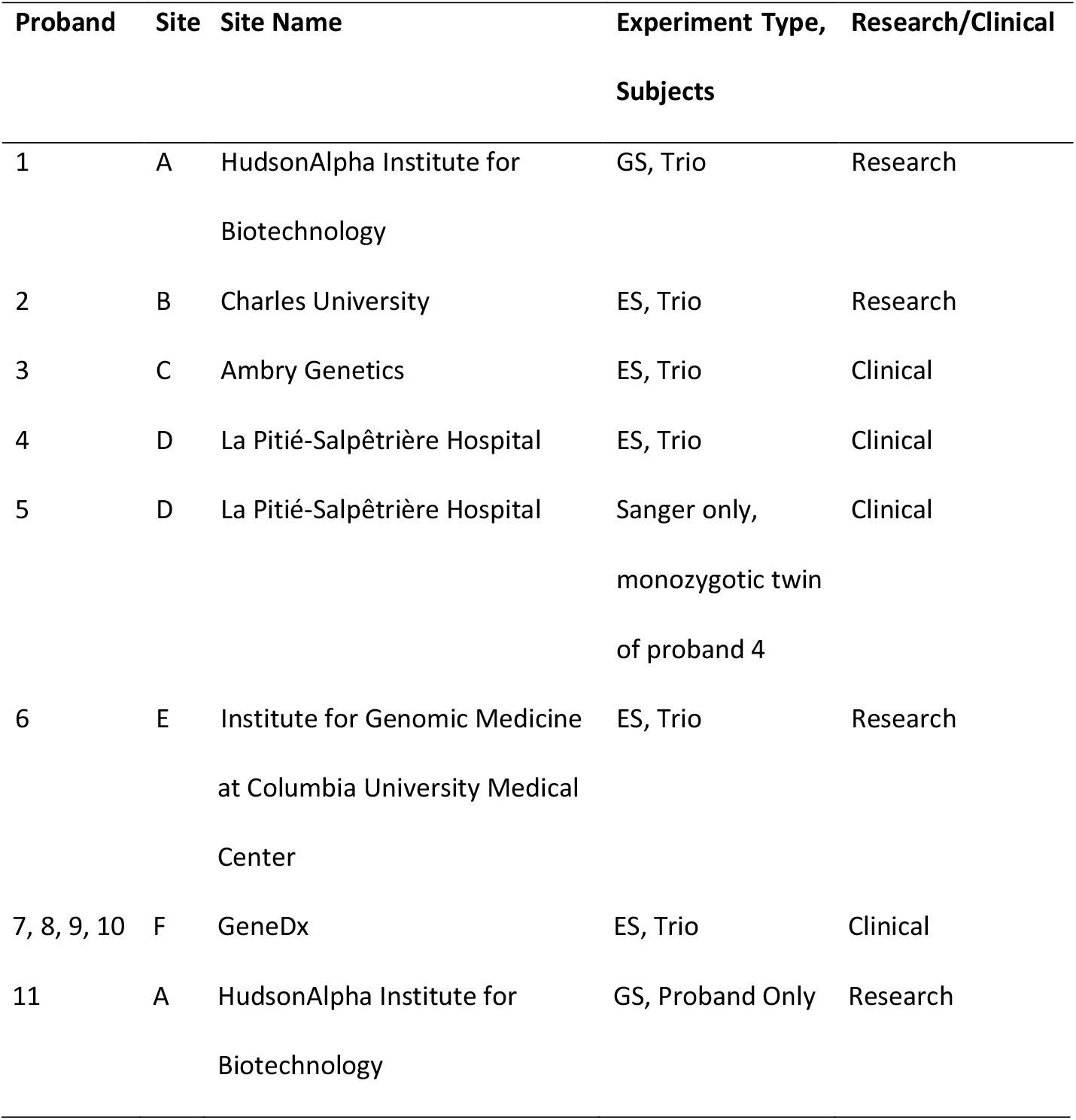

### Three dimensional modeling

The protein structure determined by Holbourn et al. [37] was used for the assessment of the potential effect of the mutations on RALA activity (PDB ID: 2BOV). The structure was visualized using PyMOL 0.99rc6[38]. Additional protein modeling was performed as previously described [39]. The GTP/GDP-binding residues of RALA were defined as those in which any atom of a residue (side chain or backbone) lies within 1.5 angstroms of an atom of the ligand.

### Cloning, protein expression, and purification

RALA cDNA was synthesized (Integrated DNA Technologies, Skokie, IL, USA) based on the coding sequence of NM_005402.3, with substitutions identified in patients described here (probands 1-9, 11; see Note below) used to represent variation. Following PCR amplification, coding sequences were cloned into Champion™ pET302/NT-His (ThermoFisher Scientific, Waltham, MA, USA, # K630203) using Gibson Assembly Master Mix (New England BioLabs, Ipswich, MA, USA, #E2611). All *RALA* coding sequences were Sanger sequenced and compared to NM_005402.3. The only differences within the coding regions of *RALA* were those observed in the probands. Single Step (KRX) Competent Cells (#L3002, Promega Corporation, Madison WI, USA) were transformed with plasmids, and bacteria were grown overnight at 37°C in LB plus ampicillin. Bacteria were diluted 1:100 in fresh LB plus 0.05% glucose and 0.1% rhamnose to induce a 6-His-tagged recombinant RALA protein. Bacteria were collected after 8 h incubation at 25°C, and snap-frozen on dry ice. 6-His-tagged proteins were purified using Dynabeads™ His-Tag Isolation and Pulldown (#10103D, ThermoFisher Scientific, Waltham, MA, USA) according to the manufacturer’s protocol. Protein purity was assessed using standard SDS-PAGE and Coomassie Blue staining. Protein concentration was quantified using a Take3 microplate reader (BioTek, Winooski, VT, USA) by assessing absorbance at 280 nm. Protein amounts were normalized among samples in Dynabead elution buffer prior to use in assays.

### GTPase activity

GTPase activity of 0.95 μg of purified, recombinant proteins was assessed using the GTPase-Glo™ Assay (#V7681, Promega Corporation, Madison WI, USA). Luminescence was quantified using an LMax II 384 Microplate Reader (Molecular Devices, San Jose, CA, USA).

### G-LISA™

Binding of purified, recombinant proteins to a proprietary Ral effector protein was assessed using the RalA G-LISA™ Activation Assay Kit (#BK129, Cytoskeleton, Inc. Denver, CO), as per the manufacturer’s protocol. Briefly, purified RALA protein was incubated in the presence or absence of 15 μM GTP (#P115A, Promega) for 1.5 h at 25°C, then 23.75 ng of purified RALA/GTP mixture was applied to the Ral-BP binding plate. A Take3 microplate reader was used for quantification of this colorimetric assay.

### Western Blot

Purified proteins were detected using a polyclonal RALA Antibody (#3526S, Cell Signaling Technology, Danvers, MA, USA) at a dilution of 1:1000, and an anti-rabbit IgG secondary antibody (#926-32211, IRDye^®^ 800CW Goat anti-Rabbit IgG, Li-cor, Lincoln, NB, USA) at a dilution of 1:20,000. An Odyssey CLx Imaging System (Li-cor, Lincoln, NB, USA) was used to visualize the Western. Relative quantification of the image was performed using Image J (https://imagej.net/).

We note that while we attempted to study the effects of all variation observed here, Proband 10 was identified after functional validation began, and the recombinant protein with the K128R variant (observed in probands 6 and 7) was not able to be expressed and purified consistently. Thus GTPase and G-LISA™ results are not shown for K128R or A158del.

## Acknowledgements

We thank all families involved in the study. This work was supported by the following grants: The National Human Genome Research Institute grant (UM1HG007301, SMH, MBN, RCR, AAH, RMM, GSB, GMC); A grant from the State of Alabama (SMH, ACEH, RMM, GSB, GMC); Ministry of Health of the Czech Republic, Grant/Award Numbers: 17-29423A, 00064203 (MHan, DP, MHav, VS, ZS); Ministry of Education of the Czech Republic, Grant/Award Number: LM2015091 (MHan, DP, MHav, VS, ZS).

---

**Figure 1. Facial features of individuals with variation in *RALA*.** Overlapping features include a broad, prominent forehead, horizontal eyebrows, epicanthus, mild ptosis, slightly anteverted nares, wide nasal bridge, short philtrum, thin upper lip vermillion with an exaggerated Cupid’s bow, pointed chin, and low-set ears with increased posterior angulation.

## Supplemental Files

**Appendix S1.** Supplemental Materials and Methods.

**Appendix S2.** Clinical summaries.

**Appendix S3.** Supplemental Figures and Tables.

---

**Figure 1 was removed because it contains identifiable images. See Page 11 for descriptions of overlapping dysmorphic features, in addition to clinical summaries in Appendix S2.**

